# Chromosome-level genome assembly of *Falcataria falcata* provides insights into ancient polyploidy and duplicate gene evolution in Fabaceae

**DOI:** 10.64898/2026.10.15.759361

**Authors:** Masaki J. Kobayashi, Osamu Chigira, Widiyatno, Mohammad Na’iem, Naoki Tani

**Affiliations:** Forestry Division, Japan International Research Centre for Agricultural Sciences, Ohwashi, Tsukuba, Ibaraki 305-8686, Japan; Kyushu Regional Breeding Office, Forest Tree Breeding Center, Forestry and Forest Products Research Institute, Forest Research and Management Organization, 2320-5 Suya, Koshi, Kumamoto 861-1102, Japan; Faculty of Forestry, Universitas Gadjah Mada, Jl. Agro Bulaksumur, Yogyakarta 55281, Indonesia; Institute of Life and Environmental Sciences, University of Tsukuba, Tennodai, Tsukuba, Ibaraki 305-8577, Japan

**Keywords:** Genome, Comparative genomics, Caesalpinioideae, Whole-genome duplication, diploidization, gene duplication

## Abstract

Whole-genome duplication (WGD) has played a fundamental role in shaping plant genome evolution, yet the timing and independence of WGD events in major Fabaceae subfamilies, including Caesalpinioideae and Papilionoideae, remain debated. Here, we present a chromosome-level genome assembly of the tropical mimosoid tree *Falcataria falcata* within Caesalpinioideae and investigate the evolutionary consequences of ancient polyploidy. The assembled genome spans 614.9 Mb with 13 chromosome-level scaffolds and 99.6% completeness based on BUSCO assessment, providing a high-quality resource for comparative genomics. Synteny analyses revealed conserved macrosyntenic structures shared across Caesalpinioideae, consistent with an ancestral WGD event predating lineage diversification. In contrast, analyses of synonymous substitution rates (Ks) and phylogenetic relationships of conserved duplicate genes support independent WGD events in Caesalpinioideae and Papilionoideae. We further show that lineage-specific variation in synonymous substitution rates is negatively associated with the retention of WGD-derived duplicates following the ancestral WGD in Caesalpinioideae, suggesting that evolutionary rate heterogeneity plays a key role in post-WGD genome evolution. In addition, recent gene duplicates that arose independently in each lineage after WGD exhibit elevated Ka/Ks ratios and are predominantly associated with defense-related functions to diverse biotic stresses, suggesting more rapid sequence evolution and functional divergence compared with WGD-derived duplicates. Together, our results reveal how ancient WGD, lineage-specific evolutionary rates, and subsequent gene duplication have jointly structured genome evolution in Fabaceae. The genome of *F. falcata* provides a valuable resource for understanding the long-term evolutionary consequences of polyploidy and the diversification of ecological adaptations in legumes.

## Introduction

Whole-genome duplication (WGD) has played a central role in the evolutionary history of plants, contributing to increased genomic complexity, functional innovation, and ecological diversification (Van de Peer et al. 2021). Recurrent polyploidization events have occurred throughout angiosperm evolution and are typically followed by diploidization through the loss of duplicated genes, subfunctionalization, and neofunctionalization of duplicated genes (Conant et al. 2014). These processes have profoundly shaped extant plant genomes and are widely recognized as drivers of long-term evolutionary success.

Fabaceae (legumes), the third–largest angiosperm family, comprises approximately 19,500 species distributed across multiple subfamilies (The Legume Phylogeny Working Group et al. 2013). Phylogenomic studies have suggested that legumes experienced one or more ancient WGD events; however, the number, timing, and phylogenetic placement of these events remain controversial (Cannon et al. 2015; Koenen et al. 2021; Zhao et al. 2021). In particular, it has been debated whether Caesalpinioideae and Papilionoideae share a single ancestral WGD or instead underwent independent WGDs following their divergence (Chen et al. 2024b; Massaro et al. 2024; McLay et al. 2022; Zhang et al. 2025).

Accurate inference of ancient WGDs is especially challenging in Fabaceae because evolutionary rates vary substantially among lineages (Chen et al. 2024b; McLay et al. 2022). Synonymous substitution rate (Ks)–based approaches are widely used to infer the timing of genome duplication, but lineage-specific variation in mutation rates can obscure or distort Ks signals, leading to conflicting interpretations (Lanfear et al. 2013; Qiao et al. 2019; Smith and Donoghue 2008). Consequently, integrative approaches that combine synteny, Ks distributions, and phylogenetic analyses of duplicated genes are increasingly required to robustly reconstruct WGD histories (Chen et al. 2024a; Ng et al. 2021).

Beyond identifying WGD events, understanding the evolutionary fate of duplicated genes is essential for elucidating how polyploidy contributes to long-term adaptation (Qiao et al. 2019). WGD-derived duplicates are often preferentially retained due to dosage balance constraints, as predicted by the gene balance hypothesis (Birchler and Veitia 2012; Conant et al. 2014). In addition, neofunctionalization, the acquisition of novel adaptive functions, is also an important mechanism to retain duplicates (Ohno 1970; Panchy et al. 2016).

*Falcataria falcata* (syn. *Albizia falcata*, *Paraserianthes falcataria*, *Falcataria moluccana*) is a fast-growing mimosoid tree species within Caesalpinioideae widely distributed in Southeast Asia and the Pacific region (Cipta et al. 2017; Hughes et al. 2025). The species has undergone repeated taxonomic reclassifications, reflecting long-standing uncertainty in the generic delimitation of mimosoid legumes (Azani et al. 2017; Barneby and Grimes 1996; Ringelberg et al. 2022). Members of the mimosoid clade are characterized by distinctive floral architectures, typically bearing actinomorphic flowers aggregated into globose or spicate inflorescences with numerous exserted stamens, valvate or imbricate perianth aestivation, and frequent heteromorphy within inflorescences (Gonçalves et al. 2024; Koenen et al. 2020). Owing to these striking floral traits, mimosoids were historically treated as a distinct subfamily (Mimosoideae), a classification that persisted until recent phylogenomic analyses demonstrated their nested placement within Caesalpinioideae (Azani et al. 2017; The Legume Phylogeny Working Group et al. 2013).

As a representative of this clade, *F. falcata* occupies a key phylogenetic position for investigating genome evolution following the ancestral WGD shared across Caesalpinioideae. Ecologically, *F. falcata* is exposed to strong biotic pressures, including severe gall rust disease caused by *Uromycladium falcatariae*, which has caused widespread damage in plantations across Southeast Asia (Doungsa-ard et al. 2015; Lelana et al. 2024). Despite its ecological and economic importance, genomic resources for *F. falcata* have remained limited, constraining detailed comparative genomic and evolutionary analyses within Fabaceae.

In this study, we present a chromosome-level genome assembly of *Falcataria falcata* and conduct comparative genomic analyses across Fabaceae. Specifically, we (i) clarify the timing and independence of WGD events in Caesalpinioideae and Papilionoideae using integrated synteny-, Ks-, and phylogeny-based approaches, (ii) examine how lineage-specific evolutionary rates shape WGD signatures and the long-term retention of duplicated genes, and (iii) contrast the evolutionary trajectories of WGD-derived and recent duplicates. Together, our results provide new insights into how ancient polyploidy and subsequent gene duplication have jointly shaped genome evolution and ecological adaptation across Fabaceae.

## Materials and methods

### Plant material, DNA extraction, genome sequencing, and assembly

Young leaf tissue was collected from a *Falcataria falcata* tree grown in a 34-year-old forest stand at Iriomote Tropical Forest Tree Breeding Technical Garden (N 24°19’44.1", E 123°54’36.6"). The leaf tissue was frozen and transferred to Dovetail Genomics (now Cantata Bio, Scotts Valley, CA, USA). An Omni-C library was constructed from the leaf tissue by Dovetail Genomics as a commercial service. A long-read sequencing library was constructed using the SMRTbell Express Template Prep Kit 2.0 (PacBio, Menlo Park, CA, USA) and sequenced on the PacBio Sequel II platform. High-fidelity (HiFi) reads were assembled using Hifiasm (Cheng et al. 2021). A Dovetail Omni-C library was prepared following the manufacturer’s protocol (Cantata Bio, Scotts Valley, CA, USA) and sequenced on the Illumina HiSeq X platform. For scaffolding, the draft assembly generated by Hifiasm and the Omni-C reads were used as input for the HiRise pipeline (Putnam et al. 2016). Assembly quality was assessed using Benchmarking Universal Single-Copy Orthologs (BUSCO) v6.0.0 with the Embryophyta odb12 dataset (Tegenfeldt et al. 2025).

### Genome annotation

The *Falcataria falcata* genome was soft-masked using RepeatMasker v2.0.2 (Chen 2004) with a custom repeat library generated by RepeatModeler v2.0.2 (Flynn et al. 2020). Protein-coding genes were predicted on the soft-masked genome using BRAKER2 v2.1.2 (Brůna et al. 2021), incorporating Viridiplantae protein sequences obtained from OrthoDB v10.1 (Kriventseva et al. 2019). The predicted genes were functionally annotated based on the protein sequence similarity to *Arabidopsis thaliana* and *Medicago truncatula* proteins using DIAMOND v2.0.15 (Buchfink et al. 2021) with an E-value cutoff of 1.0 × 10⁻¹⁰. When a *F. falcata* protein matched both *A. thaliana* and *M. truncatula* proteins, the *A. thaliana* annotation was preferentially adopted.

### Visualization of genome features and synteny

Genome features of *F. falcata*, including telomere positions, gene density, GC content, and syntenic blocks, were visualized using Circos v0.69-8 (Krzywinski et al. 2009). The canonical plant telomeric repeat motif (TTTAGGG; (Shakirov et al. 2022)) was identified using tidk v0.2.65 (Brown et al. 2025). Gene density and GC content were calculated in consecutive non-overlapping 100-kb windows using BEDTools v2.30.0 (Quinlan and Hall 2010). Macrosyntenic blocks within the *F. falcata* genome were identified by detecting closest paralogous protein pairs using DIAMOND (E-value < 1.0 × 10⁻¹⁰), followed by synteny analysis with MCScanX (Wang et al. 2012).

### Comparative genomic analysis and whole-genome duplication analysis

Single-copy orthologous genes among 17 Fabaceae species (Table 1) were identified using OrthoFinder v2.5.2 (Emms and Kelly 2019). Orthologous protein sequences were aligned using MAFFT v7.508 (Kuraku et al. 2013), and poorly aligned regions were trimmed with trimAl v1.4.rev22 (Capella-Gutiérrez et al. 2009). Maximum likelihood phylogenetic trees were inferred using IQ-TREE2 v2.2.0.3 (Minh et al. 2020).

**Table 1:** 17 Fabaceae species used for the comparative genomic analysis.

| Subfamily | Species | Reference |
| --- | --- | --- |
| Caesalpinioideae | *Acacia confusa* | Chen et al. (2024b) |
|  | *Acacia crassicarpa* | Massaro et al. (2024) |
|  | *Acacia pycnantha* | McLay et al. (2022) |
|  | *Albizia julibrissin* | Chen et al. (2024b) |
|  | *Biancaea sappan* | Chen et al. (2024b) |
|  | *Chamaecrista fasciculata* | Lee et al. (2024) |
|  | *Delonix regia* | Chen et al. (2024b) |
|  | *Entada phaseoloides* | Lin et al. (2022) |
|  | *Falcataria falcata* | This study |
|  | *Faidherbia albida* | Chang et al. (2019) |
|  | *Gleditsia sinensis* | Chen et al. (2024b) |
|  | *Mimosa bimucronata* | Chen et al. (2024b) |
|  | *Prosopis cineraria* | Sudalaimuthuasari et al. (2022) |
|  | *Senna tora* | Zhu et al. (2025) |
| Papilionoideae | *Lotus japonicus* | Sato et al. (2008) |
|  | *Medicago truncatula* | Young et al. (2011) |
|  | *Vigna angularis* | Li et al. (2024) |

Macrosyntenic relationships among species were identified by detecting orthologous proteins with DIAMOND and syntenic regions with MCScanX. Synteny visualizations were generated using SynVisio (Bandi and Gutwin 2020).

To investigate the timing of whole-genome duplication (WGD) events in Fabaceae, synonymous substitution rates (Ks) were calculated for orthologous gene pairs and duplicated paralogous gene pairs in six species: *Falcataria falcata*, *Mimosa bimucronata*, *Biancaea sappan*, *Gleditsia sinensis*, *Medicago truncatula*, and *Vigna angularis*. Orthologous gene pairs were identified based on interspecific synteny using DIAMOND and MCScanX. Within each species, the closest duplicated paralogous gene pairs located in syntenic blocks were defined as WGD-derived duplicates. Protein sequences were aligned using MAFFT, and Ks values were estimated using the yn00 program implemented in the PAML package (Yang and Nielsen 2000).

To identify WGD-derived duplicate gene pairs commonly retained across five Fabaceae species (*F. falcata*, *B. sappan*, *G. sinensis*, *M. truncatula*, and *V. angularis*), WGD-derived duplicates in *F. falcata* were divided into two sets corresponding to each paralog. Only cases in which orthologs from both sets were present in all examined species were retained. Phylogenetic relationships among these conserved WGD-derived duplicates were inferred using IQ-TREE2 following the same alignment and trimming procedures described above.

### Relationship between Ks and retained WGD-derived duplicates

The relationship between the mean Ks values of WGD-derived duplicates and the number of retained WGD-derived duplicates was examined across 11 Caesalpinioideae species with chromosome-level genome assemblies (*Acacia crassicarpa*, *A. confusa*, *F. falcata*, *Albizia julibrissin*, *Mi. bimucronata*, *Entada phaseoloides*, *Delonix regia*, *B. sappan*, *Chamaecrista fasciculata*, *Senna tora*, and *G. sinensis*). Ks values were estimated as described above. Because the number of retained WGD-derived duplicates is a count variable exhibiting overdispersion, a negative binomial generalized linear model with a log link function was fitted using the glm.nb function in the MASS package in R (version 4.1.0). Statistical significance was assessed using Wald tests.

### Functional analysis of retained WGD-derived duplicates

WGD-derived duplicate genes commonly retained across 11 Caesalpinioideae species and across mimosoid species (*A. crassicarpa*, *A. confusa*, *F. falcata*, *Al. julibrissin*, *Mi. bimucronata*, and *E. phaseoloides*) were identified using the same synteny-based approach described above. Gene Ontology (GO) enrichment analysis was performed using the topGO package v2.44.0 in R (Alexa et al. 2006), based on GO annotations of *A. thaliana* and *M. truncatula* orthologs. The top 10 significantly enriched GO terms identified using the elim algorithm were retained for downstream analyses (*P*-value < 0.05).

### Comparisons between WGD-derived duplicates and recent duplicates

In *F. falcata* and *G. sinensis*, recent duplicate gene pairs were identified as closest paralogs lacking syntenic relationships and exhibiting Ks values less than 0.2. This threshold was chosen to exclude ancient duplicates associated with WGD events and to focus on lineage-specific small-scale duplications. To compare evolutionary characteristics between WGD-derived duplicates and recent duplicates, protein sequences were aligned using MAFFT, and nonsynonymous-to-synonymous substitution rate ratios (Ka/Ks) were estimated using the yn00 program in the PAML package. To compare the distributions of Ka/Ks for the WGD-derived duplicates and recent duplicates, Wilcoxon rank-sum tests were conducted using the function, wilcox.test in R. To check whether the proportions of Ka/Ks > 1 were different between the WGD-derived duplicates and recent duplicates, Fisher’s exact tests were performed using the function, fisher.test in R. Functional enrichment analysis of recent duplicates was conducted using the same GO enrichment procedure described above.

## Results

### Assembly and characterization of the *Falcataria falcata* genome

We assembled the *Falcataria falcata* genome into 91 scaffolds totaling 614.9 Mb, with an N50 of 47.4 Mb (Table 2). Thirteen scaffolds exceeded 30 Mb, collectively accounting for 99.3% (∼610.4 Mb) of the genome, while the remaining 78 scaffolds were substantially shorter (largest ∼1.2 Mb; total ∼4.5 Mb) (Figure 1b and Table 2). Because cytogenetic analysis in *F. falcata* reports a haploid chromosome number of n = 13 (Ghosh et al. 2010), we interpreted the 13 largest scaffolds as representing chromosome-level sequences, labeled Chr1–Chr13 for downstream analyses. BUSCO assessment using the Embryophyta odb12 dataset revealed 99.6% completeness for both the full assembly and the 13-chromosome subset (Table 2), a level comparable to chromosome-level assemblies in other mimosoid species, e.g., *Acacia crassicarpa*, 99.4% completeness; *A. confusa*, 99.2%; *Albizia julibrissin*, 99.4%; *Mimosa bimucronata*, 99.4% (Chen et al. 2024b; Massaro et al. 2024). Furthermore, we detected canonical plant telomeric repeats (TTTAGGG) at both ends of all chromosomes except Chr8 and Chr13, indicating nearly full-length chromosome assemblies (Figure 1c).

**Figure 1.**
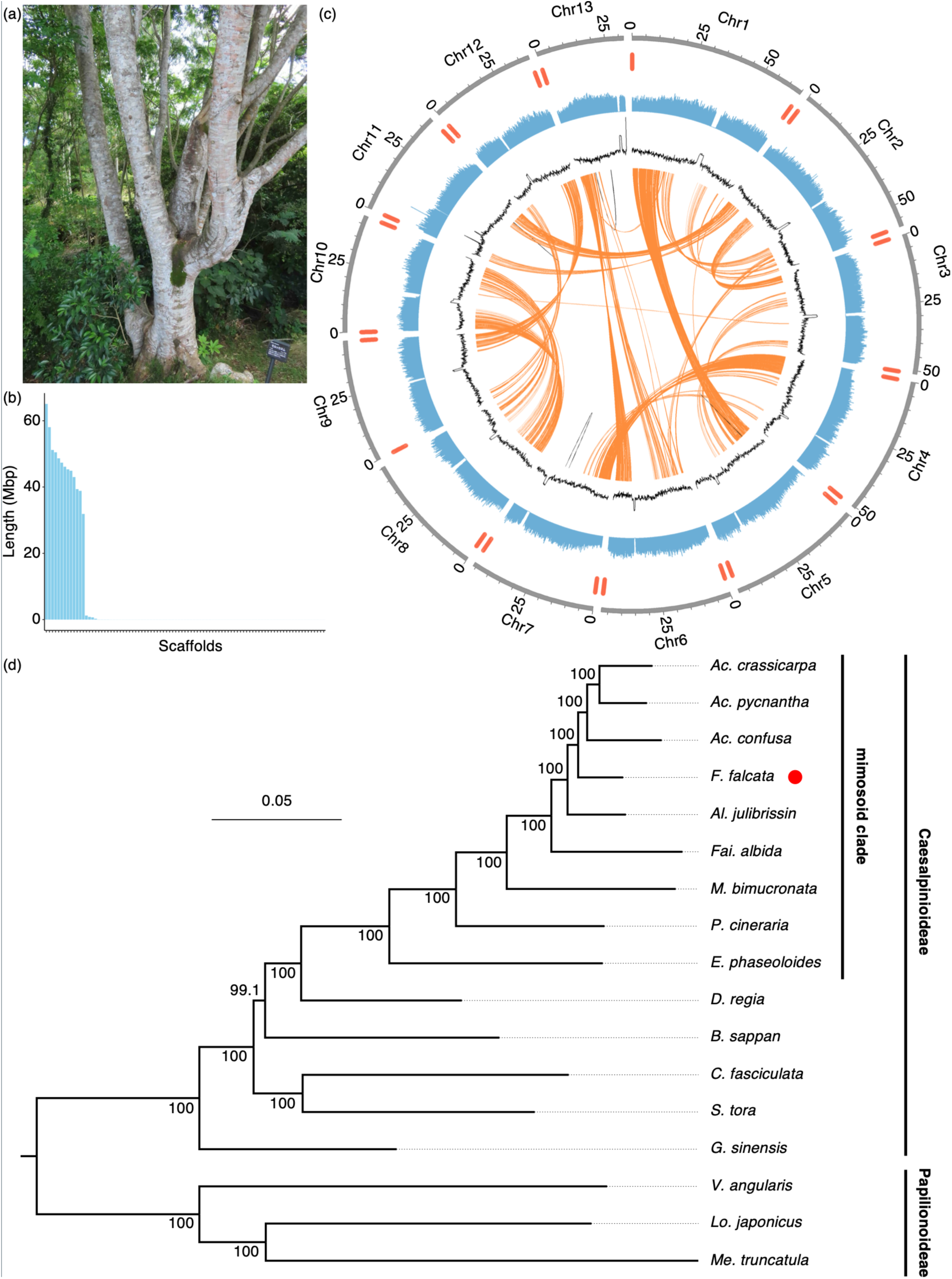
Genome assembly and phylogenetic position of *Falcataria falcata*. (a) Photograph of *Falcataria falcata*. (b) Length distribution of the 91 assembled scaffolds. The x-axis shows the 91 scaffolds ordered from longest to shortest, and scaffold lengths (Mb) are shown on the y-axis. (c) Circos plot illustrating the chromosome-level genome features of *F. falcata*. From the outermost to the innermost layers, the plot shows chromosomes, telomeric positions, gene density, GC content, and syntenic relationships. The 13 chromosomes are represented as Chr1 to Chr13. Numbers near the chromosomes indicate the positions on the chromosomes (Mb). Gene density and GC content were calculated in consecutive non-overlapping 100-kb windows. Orange and gray links represent interchromosomal and intrachromosomal synteny, respectively. (d) Maximum likelihood phylogenetic tree inferred from single-copy orthologous genes. *F. falcata* is highlighted with a red circle. The tree was constructed using IQ-TREE, and the scale bar indicates the number of substitutions per amino acid site. Node values represent ultrafast bootstrap support values. Genus abbreviations are as follows: *Ac.*, *Acacia*; *Al.*, *Albizia*; *Fai.*, *Faidherbia*; *M.*, *Mimosa*; *P.*, *Prosopis*; *E.*, *Entada*; *D.*, *Delonix*; *B.*, *Biancaea*; *C.*, *Chamaecrista*; *S.*, *Senna*; *G.*, *Gleditsia*; *V.*, *Vigna*; *Lo.*, *Lotus*; *Me.*, *Me dicago*.

**Table 2:** Statistics of *Falcataria falcata* genome assembly.

| Parameters | All scaffolds | 13 longest scaffolds |
| --- | --- | --- |
| No. of scaffolds | 91 | 13 |
| Total length of scaffolds (Mb) | 614.9 | 610.4 |
| N50 (Mb) | 47.4 | 47.4 |
| Complete BUSCOs (%) | 99.6 | 99.6 |

Gene prediction identified 44,413 protein-coding loci. This gene count is comparable to, or slightly higher than, those reported for other mimosoid species with chromosome-level genome assemblies, including *A. crassicarpa* (29,488) and *M. bimucronata* (33,062), and *Al. julibrissin* (40,866) (Chen et al. 2024b; Massaro et al. 2024). Sliding-window analysis revealed characteristic reductions in gene density concomitant with GC% peaks, suggesting the presence of a putative centromeric region in each chromosome (Figure 1c). A phylogeny constructed from 401 single-copy orthologs across 17 Fabaceae species (Fig. 1d; Table 1) recapitulated established relationships: *F. falcata* clusters closely with *Acacia* and *Albizia*, confirming the accuracy of our gene models and ortholog inference (Azani et al. 2017).

### Evidence for past WGD in *F. falcata* and timing of WGD events across Fabaceae

Synteny analysis revealed extensive conserved blocks, including Chr1–Chr5 and Chr2–Chr11, consistent with expectations for WGD events in Caesalpinioideae (Figure 1c). We defined syntenic paralogs in *F. falcata* as WGD-derived duplicates. Comparative syntenic profiling among seven Caesalpinioideae species showed near 1:1 block correspondence—even among species with differing chromosome counts—supporting a shared ancestral WGD event (Figure 2; Supplemental Figure S1).

**Figure 2.**
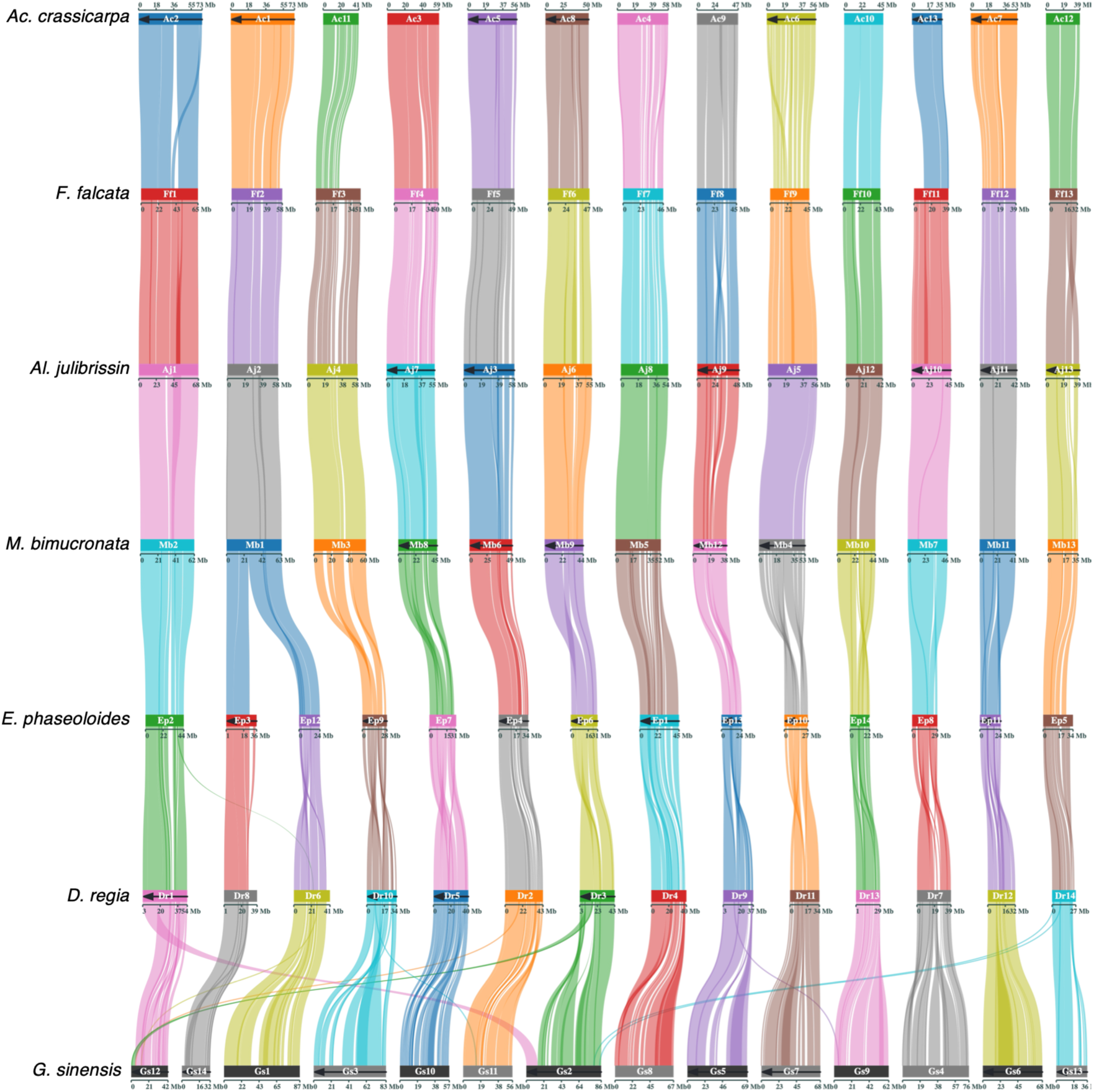
Macrosyntenic relationships among Caesalpinioideae genomes. Macrosyntenic blocks among seven species in Caesalpinioideae. Rectangles represent chromosomes, and connecting lines indicate syntenic blocks. Ten colors are used sequentially to distinguish chromosomes and syntenic regions. Left-to-right arrows along chromosomes indicate that reverse-complement sequences were used. Genus abbreviations are as follows: *Ac.*, *Acacia*; *Al.*, *Albizia*; *M.*, *Mimosa*; *E.*, *Entada*; *D.*, *Delonix*; *G.*, *Gleditsia*.

To analyze the timing of WGD events across Fabaceae, we calculated the distributions of synonymous substitution rates (Ks) for WGD-derived paralogs and interspecific orthologs in Caesalpinioideae and Papilionoideae. WGD-related Ks peaks in each examined pair in Caesalpinioideae species (*F. falcata*–*M. bimucronata*, *F. falcata*–*B. sappan*, *F. falcata*–*G. sinensis*) occurred at Ks values equal to or lower than orthologous peaks, consistent with WGD events predating species divergences (Figure 3a–c). The same pattern was observed in a pair of the Papilionoideae species, *M. truncatula*–*V. angularis* (Figure 3d). However, when comparing Caesalpinioideae with Papilionoideae (*F. falcata* vs. *M. truncatula/V. angularis*), WGD Ks peaks were higher than ortholog peaks (Figure 3e,f), indicating independent WGD events in the two subfamilies. An unrooted tree reconstructed from 70 WGD-derived gene pairs conserved across five Fabaceae species confirmed distinct Caesalpinioideae and Papilionoideae clades (Figure 3g), which further supports independent WGD events in the two subfamilies.

**Figure 3.**
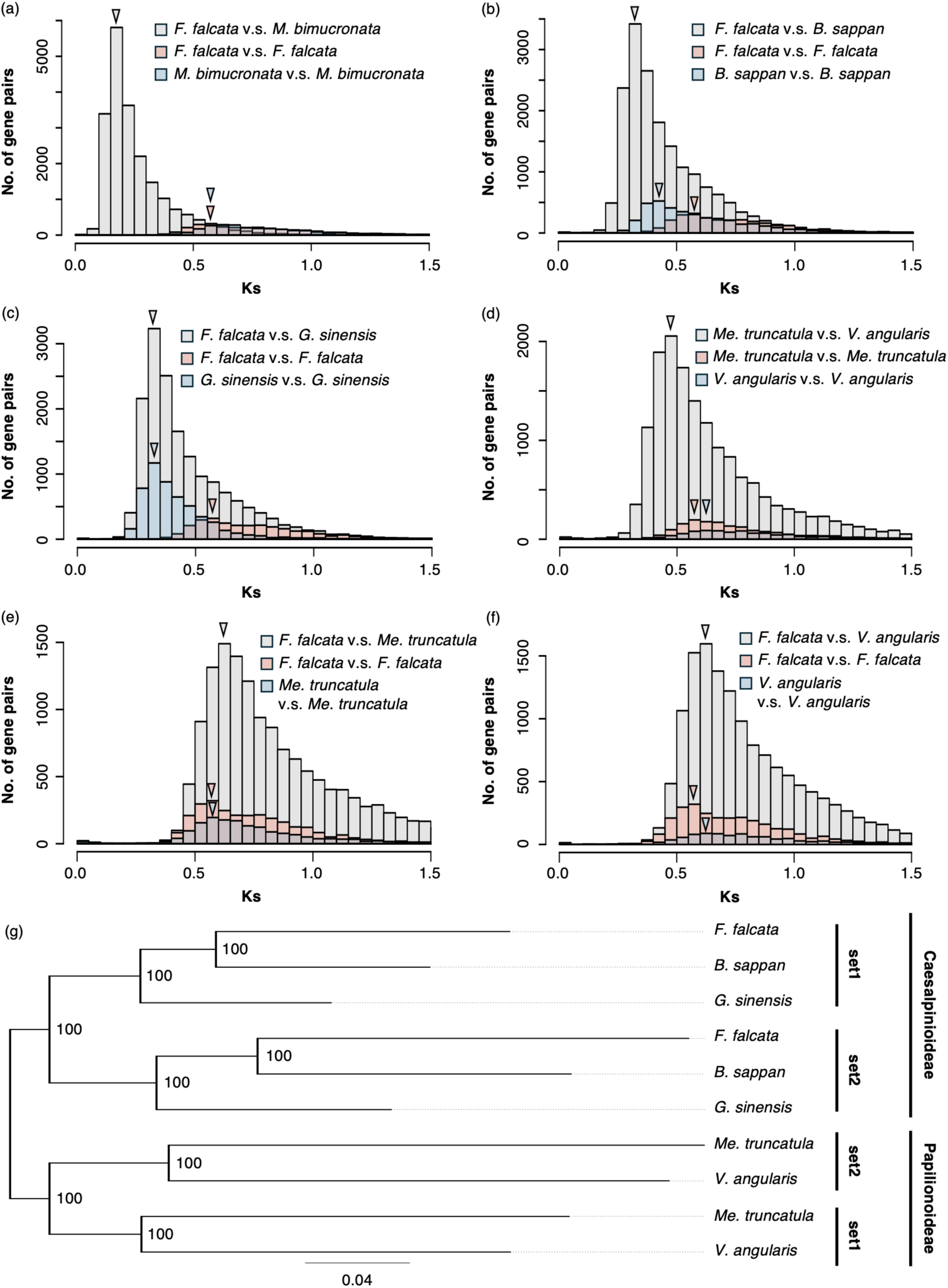
Timing of whole-genome duplication events in Caesalpinioideae and Papilionoideae. Timing of whole-genome duplication (WGD) events in Caesalpinioideae and Papilionoideae. (a–f) Distributions of synonymous substitution rates (Ks) for WGD-derived duplicate gene pairs (red and blue) and orthologous gene pairs between species (gray). Species pairs analyzed are: (a) *Falcataria falcata* and *Mimosa bimucronata*; (b) *F. falcata* and *Biancaea sappan*; (c) *F. falcata* and *Gleditsia sinensis*; (d) *Medicago truncatula* and *Vigna angularis*; (e) *F. falcata* and *M. truncatula*; and (f) *F. falcata* and *V. angularis*. Arrowheads indicate Ks peak positions. Only duplicate gene pairs with Ks values less than 1.5 are shown to improve visualization. (g) Phylogenetic relationships of WGD-derived duplicate genes commonly retained in five species: *F. falcata*, *M. bimucronata*, *B. sappan*, *G. sinensis*, *M. truncatula*, and *V. angularis*. The maximum likelihood tree was constructed using IQ-TREE, and the scale bar indicates the number of substitutions per amino acid site. Node values represent ultrafast bootstrap support values.

### Correlation between evolutionary rate and duplicate retention

Because the positions and heights of the Ks peaks differed among WGD-derived gene pairs in Caesalpinioideae species, we investigated whether variation in synonymous substitution rates (Ks) influenced retained duplicate counts across 11 Caesalpinioideae species. A negative binomial GLM revealed a significant negative correlation between mean Ks and number of retained WGD-derived duplicates (*P* = 5.0 × 10⁻⁶; Figure 4), suggesting that lineages with higher mutation rates have lower numbers of WGD-derived duplicates. *Senna tora* exhibited fewer retained duplicates than expected (Figure 4), which may be due to extensive genome rearrangements in this species (Supplemental Figure S1c).

**Figure 4.**
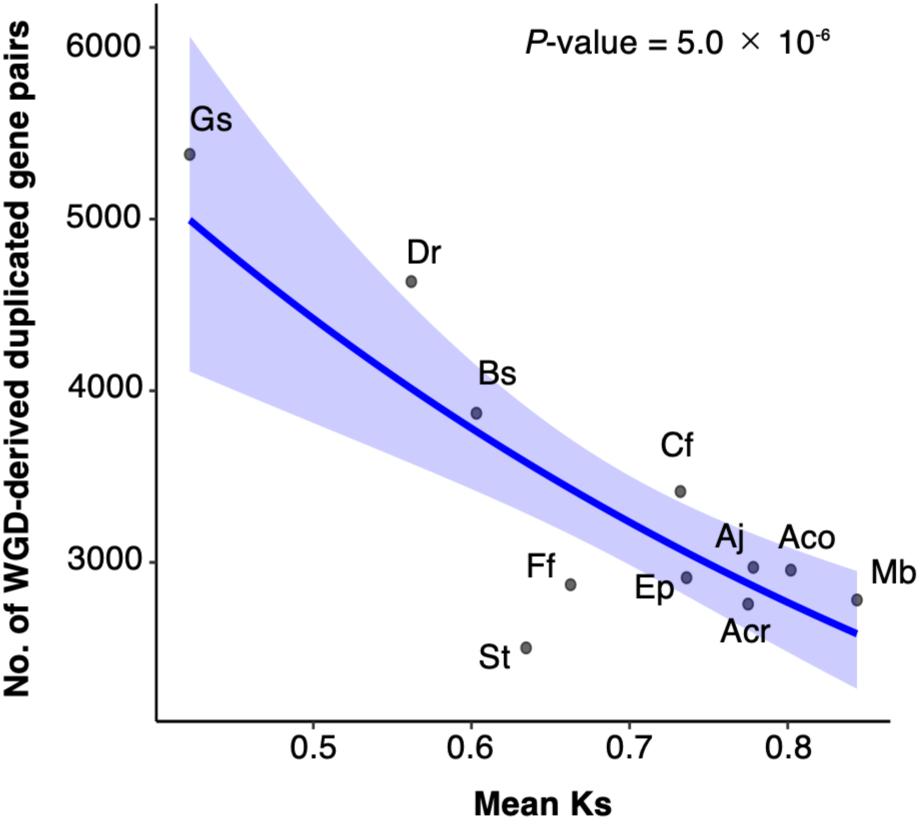
Relationship between retained WGD-derived duplicates and evolutionary distance. Relationship between the number of retained WGD-derived duplicate genes and the mean synonymous substitution rate (Ks) across 11 Caesalpinioideae species. Each point represents an individual species. The solid blue line indicates the fitted relationship estimated using a negative binomial generalized linear model with a log link function, and the shaded area represents the 95% confidence interval. A significant negative association between mean Ks values and the number of retained WGD-derived duplicates was detected (*P* = 5.0 × 10⁻⁶). Species abbreviations are as follows: Acr, *Acacia crassicarpa*; Aco, *A. confusa*; Ff, *Falcataria falcata*; Aj, *Albizia julibrissin*; Mb, *Mimosa bimucronata*; Ep, *Entada phaseoloides*; Dr, *Delonix regia*; Bs, *Biancaea sappan*; Cf, *Chamaecrista fasciculata*; St, *Senna tora*; Gs, *Gleditsia sinensis*.

### Functional properties of commonly retained WGD-derived duplicates in multiple Caesalpinioideae species

GO enrichment of 210 WGD-derived duplicates conserved across all 11 Caesalpinioideae species highlighted transcription regulation and DNA-templated transcription as overrepresented functions, in line with dosage-balance hypotheses (Birchler and Veitia 2012; Conant et al. 2014). Notably, we also found the enrichment of functions potentially related to rhizobial symbiosis and nitrogen fixation, such as cytokinin signaling, innate immunity-related processes, and nitrogen starvation response (Figure 5a). Among 1,078 WGD-derived duplicates retained specifically in the mimosoid clade, floral development and meristem patterning terms were significantly enriched (Figure 5b).

**Figure 5.**
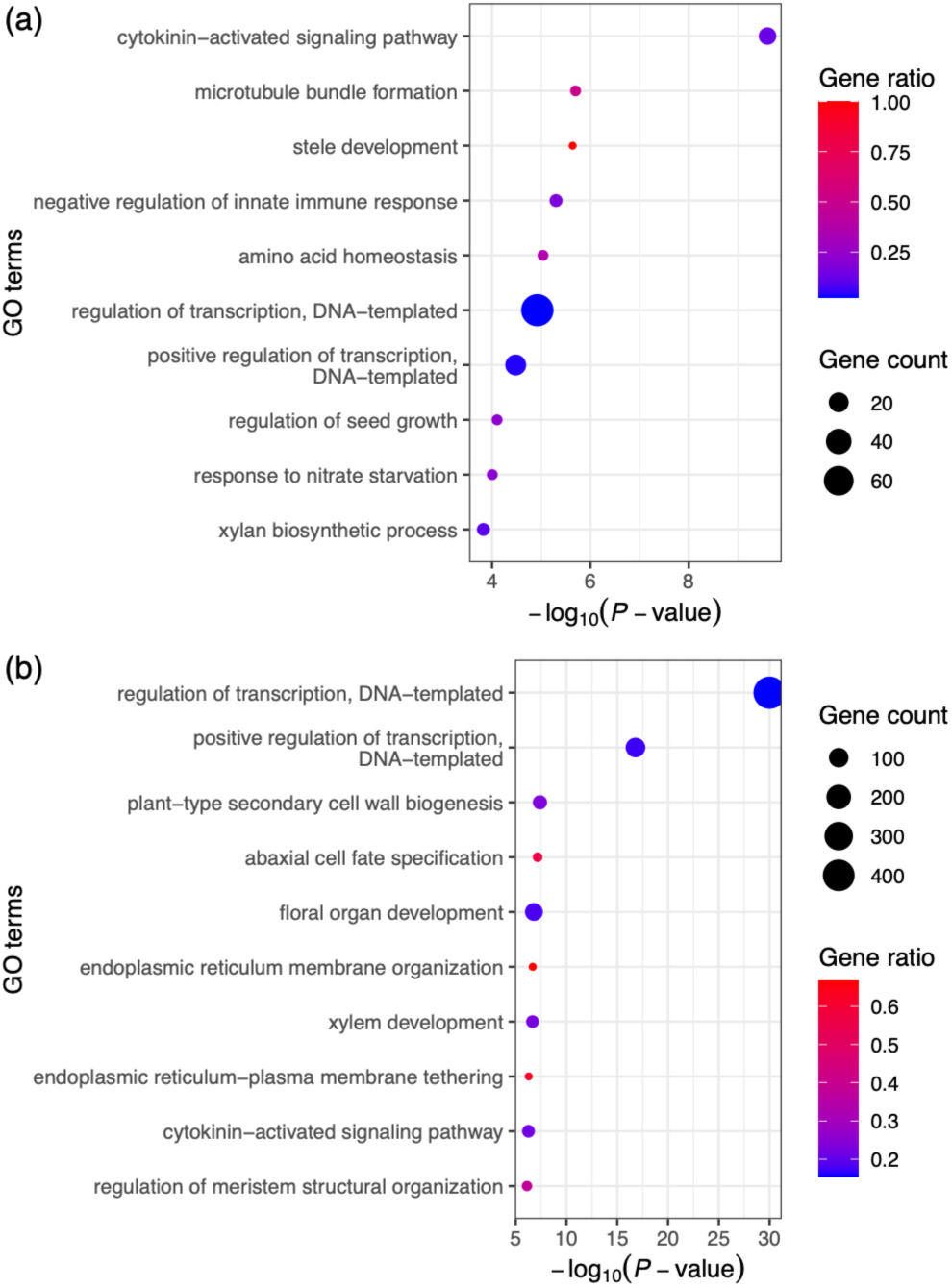
Functional enrichment of conserved WGD-derived duplicates. Gene Ontology (GO) enrichment analysis of WGD-derived duplicate genes commonly retained in Caesalpinioideae (a) and mimosoid species (b). Dot colors represent the gene ratio between retained WGD-derived duplicates and all genes in each GO category, with red indicating a higher gene ratio. Dot sizes correspond to the number of enriched genes within each GO term.

### Comparisons between WGD-derived duplicates and recent duplicates

We identified a substantial number of duplicated paralogs with small Ks values in Caesalpinioideae species and defined those with Ks < 0.2 as recent duplicates to distinguish from WGD-derived duplicates (Figure 6a,b; Supplemental Figure S2). Recent duplicates showed significantly elevated Ka/Ks distributions compared with WGD-derived paralogs in both *F. falcata* and *G. sinensis* (Wilcoxon rank-sum test, *P* < 0.05; Figure 6c,d), with many Ka/Ks values >1. In *F. falcata*, 28.0% of recent duplicate gene pairs exhibited Ka/Ks values greater than 1.0, compared with only 0.1% of WGD-derived duplicates. A similar pattern was observed in *G. sinensis*, where 13.0% of recent duplicates showed Ka/Ks > 1.0, whereas only 0.0% of WGD-derived duplicates exceeded this threshold. The difference in the proportion of gene pairs with Ka/Ks > 1.0 between recent and WGD-derived duplicates was statistically significant in both species (Fisher’s exact test, P < 0.05). GO analysis indicated enrichment for defense-related processes in both species (Figure 6e,f). However, recent duplicates in *F. falcata* implicated in fungal, bacterial, and oomycete responses (Figure 6e), whereas those in *G. sinensis* were predominantly associated with herbivore and wounding responses (Figure 6f). This may reflect lineage-specific ecological adaptations affecting recent duplicate evolution.

**Figure 6.**
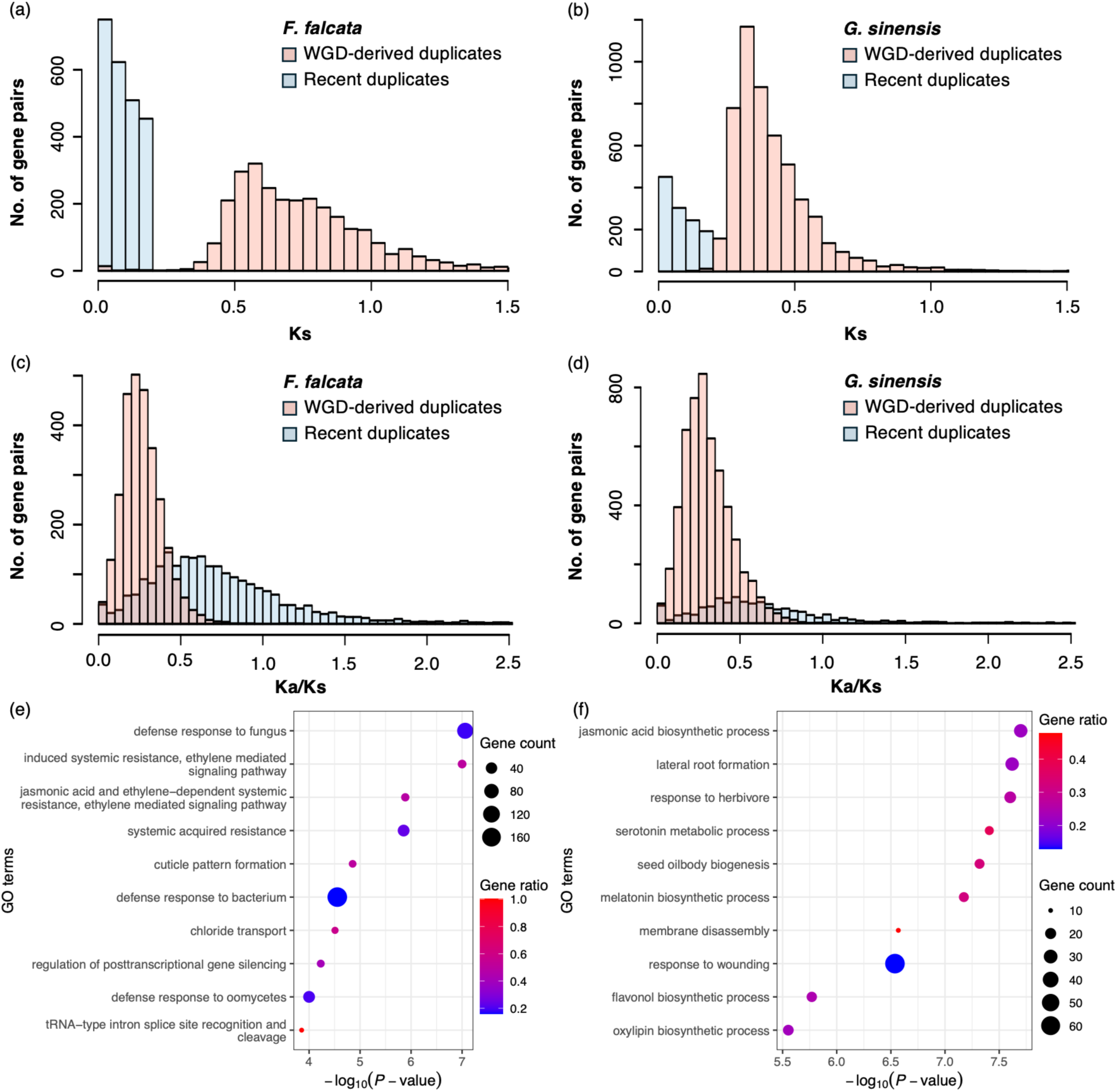
Contrasting evolutionary patterns of WGD-derived and recent duplicates. Comparisons between WGD-derived duplicate genes and recent duplicate genes. (a, b) Distributions of synonymous substitution rates (Ks) for WGD-derived duplicates (red) and recent duplicates (blue; Ks < 0.2) in *Falcataria falcata* (a) and *Gleditsia sinensis* (b). (c, d) Distributions of nonsynonymous-to-synonymous substitution rate ratios (Ka/Ks) for WGD-derived duplicates (red) and recent duplicates (blue) in *F. falcata* (c) and *G. sinensis* (d). Only duplicate gene pairs with Ks < 1.5 and Ka/Ks < 2.5 are shown to improve visualization. (e, f) GO enrichment analysis of recent duplicate genes in *F. falcata* (e) and *G. sinensis* (f). Dot colors represent gene ratios, and dot sizes indicate the number of enriched genes per GO term.

## Discussion

### Independent whole-genome duplication (WGD) events in Caesalpinioideae and Papilionoideae

The timing and phylogenetic placement of whole-genome duplication (WGD) events in Fabaceae have long been debated. Some studies have proposed independent WGDs in Caesalpinioideae and Papilionoideae (Cannon et al. 2015; Chen et al. 2024b; Massaro et al. 2024; McLay et al. 2022; Zhao et al. 2021), whereas others have suggested a single WGD event in their common ancestor (Koenen et al. 2021; Zhang et al. 2025). To address this issue, we conducted a comprehensive study incorporating synteny-, Ks-, and phylogeny-based analysis.

The chromosome-level genome assembly of *F. falcata* provides compelling evidence for an ancient whole-genome duplication (WGD) shared across Caesalpinioideae. Extensive macrosynteny observed among *F. falcata* (Fig. 1c) and multiple Caesalpinioideae species (Fig. 2), including *Acacia crassicarpa*, *Albizia julibrissin*, *Mimosa bimucronata*, *Entada phaseoloides*, *Delonix regia*, and *Gleditsia sinensis*, revealed largely one-to-one chromosomal correspondence despite variation in chromosome number (13 or 14). This pattern strongly supports a single ancestral WGD predating the diversification of extant Caesalpinioideae lineages, consistent with recent chromosome-level comparative genomic analyses of this subfamily (Chen et al. 2024b). Ks-based analyses provide independent support for this inference (Fig. 3a-c).

In contrast, comparisons between Caesalpinioideae and Papilionoideae species revealed a different evolutionary history. When *F. falcata*was compared with *Medicago truncatula* or *Vigna angularis*, orthologous Ks peaks were consistently older than WGD-derived Ks peaks within each subfamily (Figure 3e,f), indicating that species divergence preceded the respective WGD events. Moreover, phylogenetic analysis of WGD-derived duplicate genes commonly retained across Fabaceae demonstrated that Caesalpinioideae and Papilionoideae duplicates formed distinct clades (Figure 3g), providing independent phylogenetic support for separate WGD events in these two subfamilies.

These results are consistent with previous phylogenomic studies suggesting multiple, lineage-specific WGDs in Fabaceae rather than a single pan-legume polyploidization event (Cannon et al. 2015). Our findings thus refine the evolutionary framework of Fabaceae by clarifying that Caesalpinioideae and Papilionoideae underwent independent genome duplications after their divergence.

### Lineage-specific evolutionary rates shape WGD signal divergence

Although Caesalpinioideae species share a common WGD origin, the positions and shapes of Ks peaks varied markedly among species. Notably, *F. falcata* and *M. bimucronata* exhibited similar WGD-related Ks peaks (Figure 3a), whereas *Biancaea sappan* and *Gleditsia sinensis* showed substantially lower Ks values (Figure 3b,c). This discrepancy is more plausibly explained by lineage-specific differences in synonymous substitution rates than by differences in WGD timing.

This interpretation is supported by the species phylogeny (Figure 1d), in which branch lengths varied considerably among Caesalpinioideae lineages. Species belonging to the mimosoid clade tended to show longer branches, indicating higher mutation rates, whereas *Delonix regia* and *G. sinensis* exhibited shorter branches. Similar rate heterogeneity has been reported in other comparative genomic analyses of Caesalpinioideae and legumes more broadly (Chen et al. 2024b; Qiao et al. 2019). Because substitution rates in plants have been reported to vary dependently on life-history traits, such as woody versus herbaceous growth (Smith and Donoghue 2008) and plant height (Lanfear et al. 2013), the observed variation in synonymous substitution rates among Caesalpinioideae lineages may be associated with their diverged life-history traits (Azani et al. 2017).

These findings highlight an important caveat in interpreting Ks-based WGD dating: identical duplication events can produce divergent Ks peaks across lineages when mutation rates differ substantially. The highly variable mutation rates in Fabaceae complicate the inference of WGD timing based solely on Ks values and likely contributed to divergent interpretations in previous studies (Chen et al. 2024b; Zhao et al. 2021). Consequently, integrated analysis incorporating synteny-, Ks-, and phylogeny-based results remains essential for robust inference of shared WGD events.

### Evolutionary rate and duplicate gene retention after WGD

Having clarified the timing and independence of WGDs in Fabaceae, we next examined how lineage-specific evolutionary rates have influenced the long-term retention of duplicated genes following these events. Following WGD, genomes undergo progressive diploidization characterized by extensive gene loss (Conant et al. 2014; Qiao et al. 2019). Across 11 Caesalpinioideae species, we detected a significant negative association between mean Ks values of WGD-derived duplicates and the number of retained duplicates (Figure 4). This pattern suggests that lineages with higher synonymous substitution rates experience faster erosion of duplicated gene pairs. Such a relationship aligns with theoretical and empirical studies indicating that accelerated sequence evolution promotes fractionation of duplicated genes following polyploidization (Panchy et al. 2016; Qiao et al. 2019). The outlier behavior of *Senna tora*, which retained fewer duplicates than expected based on Ks alone, may reflect additional genome-scale processes, such as extensive chromosomal rearrangements (Supplemental Figure S1c), that further accelerate duplicate loss.

### Functional constraints on retained WGD-derived duplicates

Consistent with the gene dosage balance hypothesis (Birchler and Veitia 2012; Conant et al. 2014), WGD-derived duplicate genes commonly retained across Caesalpinioideae were significantly enriched for transcriptional regulation and DNA-templated transcription (Figure 5). Genes involved in regulatory networks and macromolecular complexes are thought to be preferentially retained after WGD to preserve stoichiometric balance, whereas small-scale duplications of such genes are often deleterious (Birchler and Veitia 2012; Conant et al. 2014).

Beyond these canonical categories, we detected enrichment of genes associated with cytokinin signaling, innate immune regulation, and responses to nitrogen starvation in the conserved WGD-derived duplicates in Caesalpinioideae (Figure 5a). These functions may be particularly relevant to legumes, which rely on complex signaling pathways for symbiosis with nitrogen-fixing bacteria (Yang et al. 2022). The preferential retention of such genes suggests that WGD may have contributed to the evolutionary refinement of symbiotic and nitrogen-response pathways in Caesalpinioideae.

Among mimosoid species, additional enrichment for floral organ development and meristem organization was observed (Figure 5b). Species in the mimosoid clade have a set of conserved peculiar floral characteristics, such as radial symmetry, valvate aestivation of the petals, presence of polyandry and polycarpelly (Azani et al. 2017; Koenen et al. 2020; Paulino et al. 2014; Tucker 2003), due to which mimosoid used to be ranked as a subfamily (Azani et al. 2017; The Legume Phylogeny Working Group et al. 2013). In addition to these conserved floral characteristics, they also present enormous variations in floral characteristics and high merism labilities (Gonçalves et al. 2024). Thus, this result implies that commonly retained WGD-derived duplicates may have played a role in the specialization of the floral morphologies compared with other lineages and the diversification of floral characteristics and meristem labilities within this clade.

### Contrasting evolutionary trajectories of WGD-derived and recent duplicates

In contrast to WGD-derived duplicates, recent duplicates (Ks < 0.2) in *F. falcata* and *G. sinensis* exhibited markedly higher Ka/Ks ratios, with many gene pairs showing Ka/Ks values exceeding 1.0 (Figure 6c,d). This pattern indicates relaxed constraint or positive selection, consistent with functional divergence following recent gene duplication (Yang 2019).

Functional enrichment analyses revealed that recent duplicates were predominantly associated with defense-related processes (Figure 6e,f). Interestingly, the dominant defense categories differed between species: *F. falcata* recent duplicates were enriched for responses to microbial pathogens (Figure 6e), whereas *G. sinensis* showed enrichment for herbivore- and wounding-related responses (Figure 6f).

In *F. falcata*, gall rust disease, caused by the fungal pathogen *Uromycladium falcatariae*, severely affects tree growth and mortality (Doungsa-ard et al. 2015), becoming a major threat in *F. falcata* plantations (Lelana et al. 2024). In addition, *F. falcata* is susceptible to a wide range of fungal pathogens throughout its life cycle, including damping-off (*Pythium* spp., *Phytophthora* spp. and *Rhizoctonia* spp.), anthracnose (*Colletotrichum* spp.), and root rot diseases (*Botryodiplodia* spp., *Ganoderma* spp., *Ustulina* spp., and *Rosellinia* spp.) (Nair and ed. 2000). The humid environments inhabited by *F. falcata* may increase exposure to diverse microbial antagonists, suggesting that microbial pathogens may have historically represented an important selective pressure in *F. falcata*.

In contrast, species in the genus *Gleditsia*, including *G. sinensis*, possess long thorns (Xiao et al. 2025), which are defensive organs against herbivore attack, and are especially effective against small, browsing mammals (Milewski et al. 1991). This suggests that herbivore attacks may have been a major threat for *G. sinensis*, which may explain the observed functional enrichment of herbivore-and wounding-related responses in recent duplicates of this species. Such lineage-specific functional biases suggest that recent gene duplications provide a flexible genetic substrate for adaptation to local ecological pressures.

This contrast between WGD-derived duplicates and recent duplicates supports a two-phase model of duplicate gene evolution in legumes: WGD-derived duplicates tend to retain ancestral regulatory functions under strong dosage constraints, whereas recent duplicates undergo rapid sequence divergence and contribute to adaptive innovation (Panchy et al. 2016; Qiao et al. 2019).

## Conclusions

Together, our analyses demonstrate that an ancient WGD shared across Caesalpinioideae has been differentially shaped by lineage-specific mutation rates, genome rearrangements, and selective pressures. The genome of *F. falcata* provides a valuable reference for understanding the long-term evolutionary consequences of WGD in legumes and illustrates how ancient polyploidy and subsequent gene duplication have jointly contributed to genomic diversity and ecological adaptation across Fabaceae.

## Acknowledgments

We would like to express appreciation to Mr. H. Hamamoto, for assistance with sample collection and project management for Mr. O. Shimada, Mr. T. Inamoto, Dr. E. Fukatsu, Dr. M. Takahashi, Dr. K. Hayashi. This study was primarily supported by the SATREPS project “Strengthening Tropical Forest Resilience Based on the Management and Utilization of Genetic Resources Capable of Climate Change Adaptation” (JPMJSA2101), funded by the Japan Science and Technology Agency (JST) and the Japan International Cooperation Agency (JICA), and partly supported by the Japan International Research Center for Agricultural Sciences (JIRCAS) projects “Evaluation of Genetic Resources for Strengthening Productivity and Adaptability of Tropical Forests” and “Developing Climate-Resilient Technologies for Tropical Forestry.”

## Data availability

The genome assembly and annotation of *Falcataria falcata* were deposited to Figshare (https://doi.org/10.6084/m9.figshare.32309982). The genome information of *Arabidopsis thaliana* was downloaded from TAIR (https://www.arabidopsis.org). The genome information of *Medicago truncatula* and *Chamaecrista fasciculata* were downloaded from Phytozome 14 (https://phytozome-next.jgi.doe.gov). The genome information of *Acacia crassicarpa* was downloaded from DRYAD (https://doi.org/10.5061/dryad.573n5tbdr). The genome information of *Acacia pycnantha* (GCA_025563575.1), *Prosopis cineraria* (GCF_029017545.1), *Senna tora* (GCA_014851425.1), and *Vigna angularis* (GCF_016808095.1) were downloaded from National Center for Biotechnology Information (NCBI) (https://www.ncbi.nlm.nih.gov/datasets/genome). The genome information of *Acacia confusa* (https://doi.org/10.6084/m9.figshare.25000859), *Albizia julibrissin* (https://doi.org/10.6084/m9.figshare.25037561), *Biancaea sappan* (https://doi.org/10.6084/m9.figshare.25037618), *Delonix regia* (https://doi.org/10.6084/m9.figshare.25000856), *Gleditsia sinensis* (https://doi.org/10.6084/m9.figshare.25037543), and *Mimosa bimucronata* (https://doi.org/10.6084/m9.figshare.25037582) were downloaded from Figshare. The genome information of *Faidherbia albida* was downloaded from GigaDB (https://doi.org/10.5524/101054).

The genome information of *Entada phaseoloides* was downloaded from China National GeneBank (CNGB) (https://ftp.cngb.org/pub/CNSA/data1/CNP0001900/CNS0383647/CNA0029377/). The genome information of *Lotus japonicus* was downloaded from https://www.kazusa.or.jp/lotus/index.html.

## Author contributions

The authors confirm their contributions to the manuscript as follows: MJK: study design, data analysis and interpretation, and original draft preparation; OC: plant sample management and sample collection; MN and Widiyatno: study conception and project administration; NT: study conception, project administration, and data collection. All authors reviewed the results, revised the manuscript, and approved the final version

## Competing interest

The authors declare no competing interest.

## Supplemental Figures

**Figure S1.**
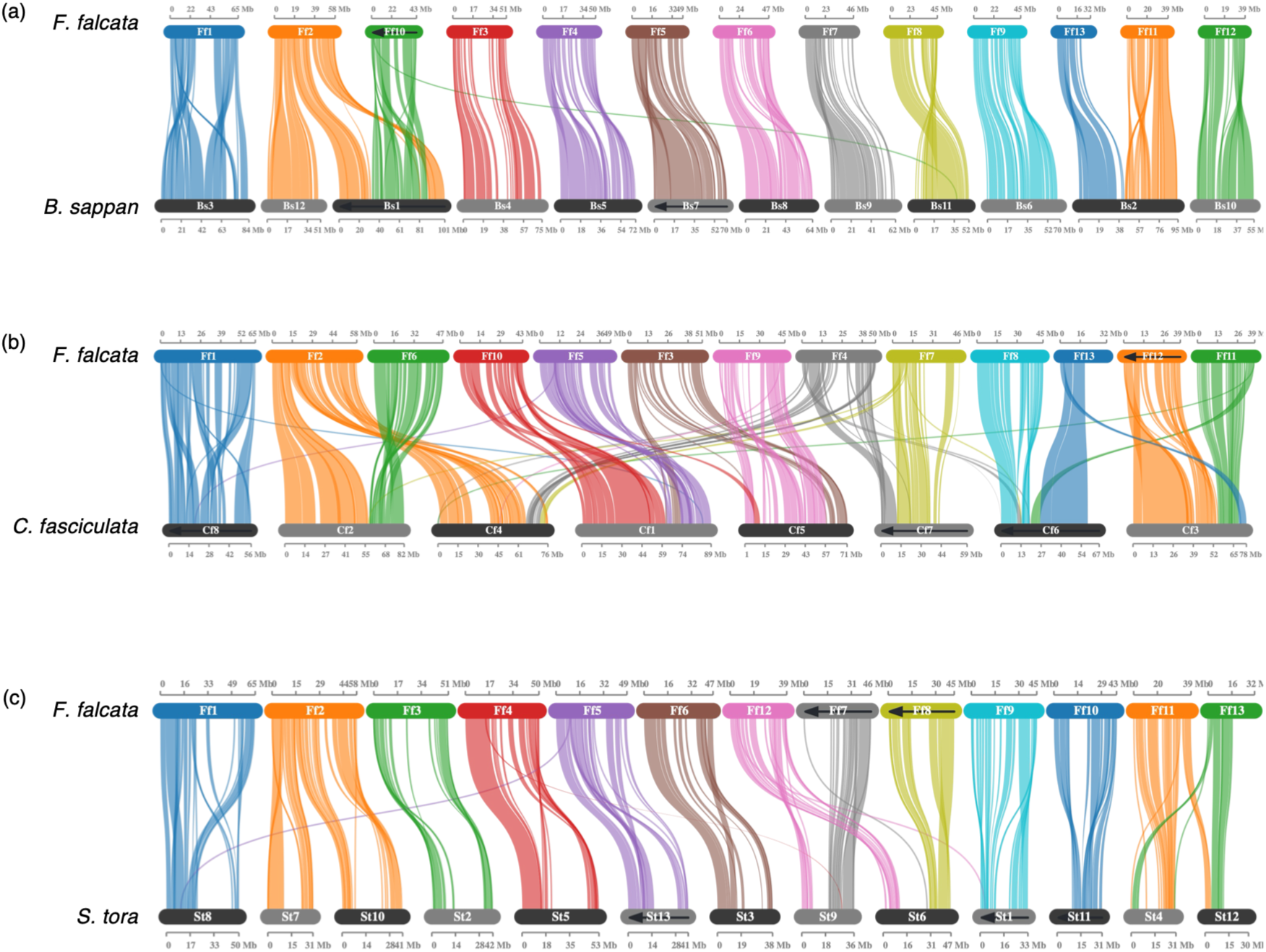
Macrosyntenic relationships between *Falcataria falcata* and other Caesalpinioideae species. Macrosyntenic blocks between *Falcataria falcata* and three Caesalpinioideae species. (a) *Biancaea sappan*, (b) *Chamaecrista fasciculata*, and (c) *Senna tora*. Rounded rectangles represent chromosomes, and connecting lines indicate syntenic blocks. Chromosomes and syntenic regions are sequentially colored using ten distinct colors. Left-to-right arrows along chromosomes indicate that reverse-complement sequences were used.

**Figure S2.**
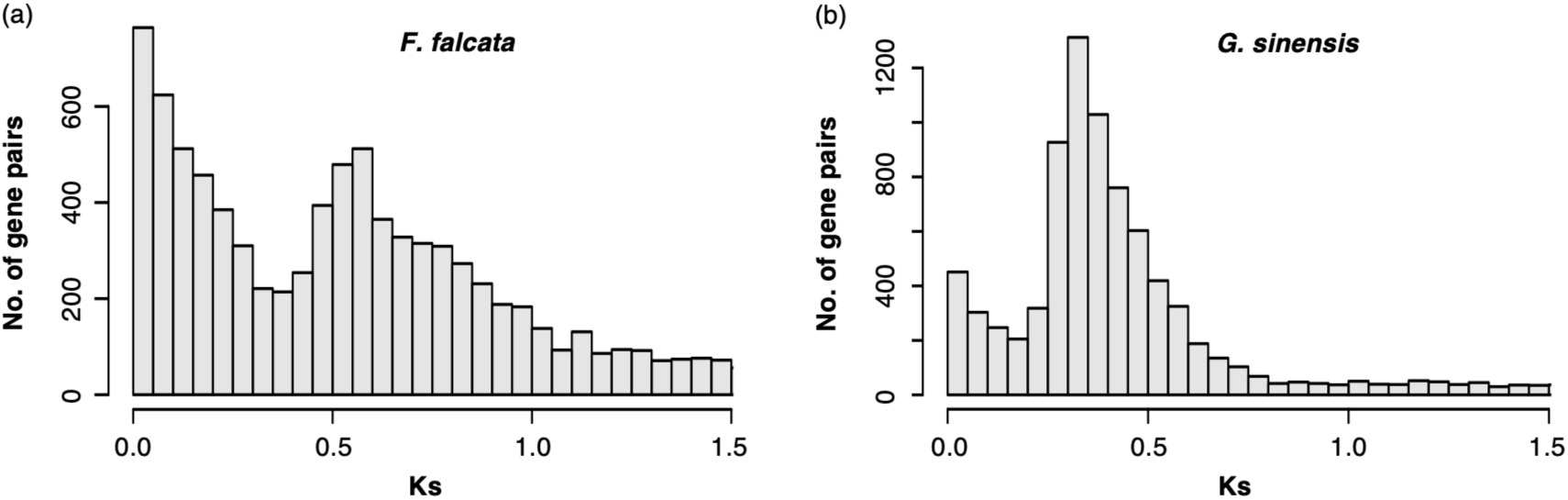
Distribution of synonymous substitution rates for duplicate genes. Distribution of synonymous substitution rates (Ks) for all duplicate gene pairs in *Falcataria falcata* (a) and *Gleditsia sinensis* (b). Only duplicate gene pairs with Ks values less than 1.5 are shown to improve visualization.

## Notes

### Competing Interest Statement

The authors have declared no competing interest.

